# A novel method for classification of tabular data using convolutional neural networks

**DOI:** 10.1101/2020.05.02.074203

**Authors:** Ljubomir Buturović, Dejan Miljković

## Abstract

Convolutional neural networks (CNNs) represent a major breakthrough in image classification. However, there has not been similar progress in applying CNNs, or neural networks of any kind, to classification of tabular data. We developed and evaluated a novel method, TAbular Convolution (TAC), for classification of such data using CNNs by transforming tabular data to images and then classifying the images using CNNs. The transformation is performed by treating each row of tabular data (i.e., vector of features) as an image filter (kernel), and applying the filter to a fixed base image. A CNN is then trained to classify the filtered images. We applied TAC to classification of gene expression data derived from blood samples of patients with bacterial or viral infections. Our results demonstrate that off-the-shelf ResNet can classify the gene expression data as accurately as the current non-CNN state-of-the-art classifiers.

## 1 Introduction

Neural networks have demonstrated tremendous progress in classification of a variety of data types. In particular, convolutional neural networks (CNNs) approach or surpass human-level accuracy for classification of images, in certain scenarios [1]. However, progress in applying neural networks for classification of tabular data has been slower. Consequently, non-neural-network methods such as XGBoost, LightGBM, Support Vector Machines (SVM) and others still dominate when using tabular data.

Several approaches have been proposed to harness the classification power of neural networks using tabular data. One of the research directions has been conversion of tabular data to images and subsequent classification of the images using the CNNs. Sharma et al. [2] pioneered an approach involving t-SNE mappings of input features (as opposed to input feature vectors). After transposing the input data matrix, they map the features on a 2D space and generate image which reflects feature similarities. The new input data points are then placed as pixels on the resulting 2D plot, and the resulting images are classified using CNN. Authors reported very good results, however using pilot and synthetic datasets of limited practical interest. An important limitation of the method is that for low-dimensional data (say, dozens or hundreds of features, which is very common in tabular data) it creates very few pixels, which may make it difficult for CNN to classify the images.

In this paper, we propose a novel method for classification of tabular data using CNNs, called *TAC* (TAbular Convolution). The core invention is a method for converting tabular data into images using convolution of a fixed *base* image and input feature vectors treated as kernels (filters). The filtered images are subsequently classified using a CNN. We demonstrate application of TAC to classify gene expression data from 2,590 blood samples from patients with an infectious disease to diagnose whether they have a bacterial or viral infection. This is a highly relevant medical problem with potentially significant impact in primary care and emergency medicine. We compared TAC accuracy for classifying the infections with the performance of advanced non-CNN algorithms. We used an off-the-shelf CNN, ResNet [3].

### Algorithm 1 Tabular Convolution (TAC). |*A*| is the cardinality of set *A*. 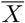 is the mean value of elements of vector *X*. * is the convolution operator

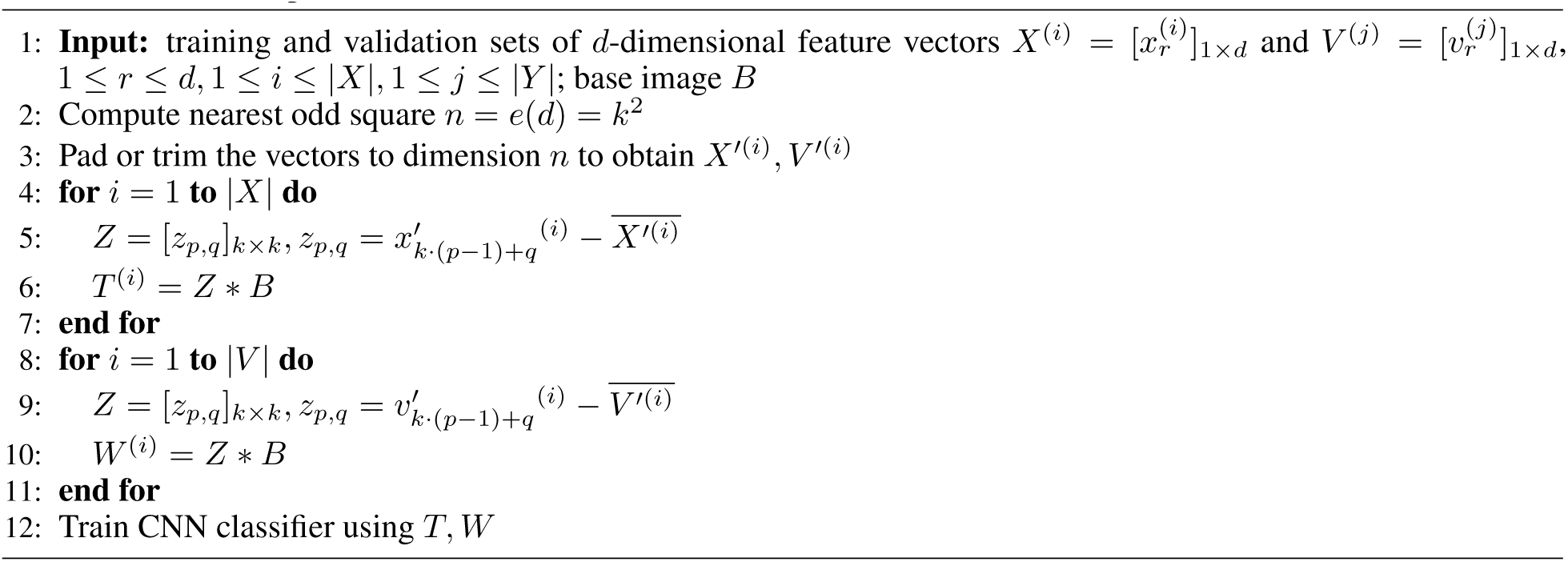

The main contributions of our work are:

- Invention of a kernel method, TAC, for conversion of tabular data to images.
- Application of the method to real-world data for classification problem of major clinical significance.
- Rigorous comparison of untuned TAC with non-CNN classifiers.
- Demonstrating that TAC classification accuracy matches non-CNN classifiers.

## 2 Methods and Materials

Our approach treats each feature vector as an image filter (kernel), convolves it with a fixed image called a *base image*, and classifies the filtered images. Therefore, TAC has the following steps (Algorithm 1):

1. Convert feature vectors to kernels.
2. Convolve the kernels with the base image.
3. Classify the filtered images using CNN.

TAC starts by converting the feature vector into a kernel by rearranging it into a square matrix, and subtracting the vector mean value. The convolutional kernels used in image processing are typically a square matrix with odd number of rows and columns (e.g., 3 × 3, 5 × 5, 7 × 7 etc). Hence, feature vectors can be directly converted to kernels if the number of features is a square of an odd integer greater than 1 (i.e., number of features must be 9, 25, 49 etc). For simplicity, we refer to these integers as *odd squares*. When the number of input features is not an odd square, the feature vectors have to be padded or trimmed to the nearest odd square. Padding can be achieved, for example, by adding zeros, random noise or engineered features (products, fractional polynomials, etc.) to the original feature vectors. Trimming is performed by removing features in an unsupervised manner, to avoid bias. For example, one could remove features with least variance. The decision of padding vs. trimming also depends on the number of features that have to be added or removed. Typically, one would identify the two nearest odd squares and trim or pad feature vectors to the nearer one. We designate the nearest odd square of *d* as *e*(*d*).

Each feature vector is zero-centered before convolution to remove the non-informative, low-frequency signal. Fig. 1 illustrates the training phase of the TAC algorithm for the case of 25-dimensional input vectors. It is assumed that feature vectors are already padded or trimmed. The CNN is trained from scratch, starting with randomly-initialized weights.

**Figure 1:**
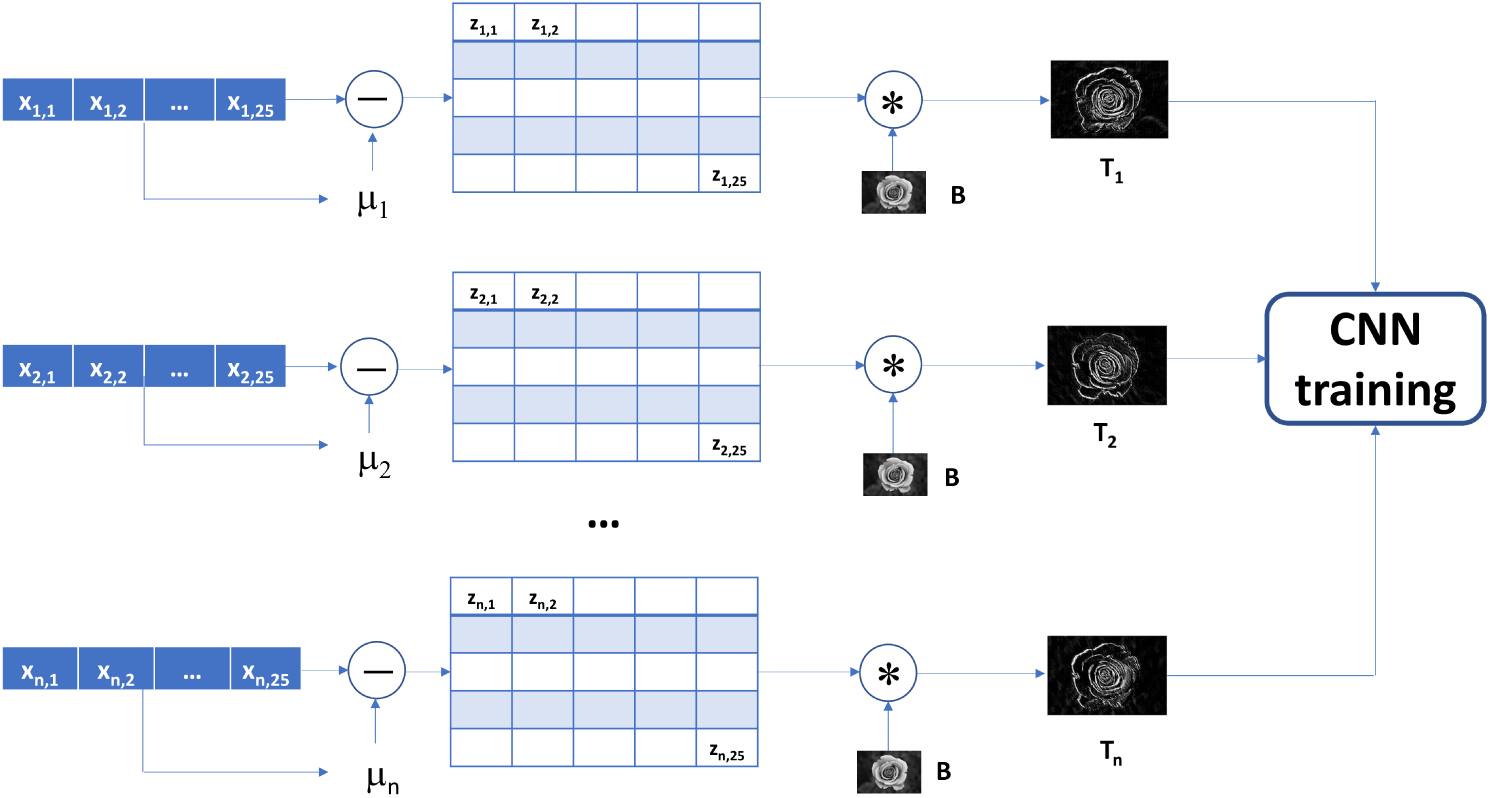
The TAC algorithm in training phase. In the example, input feature vectors have 25 elements. *µ*_*i*_ is the mean value of input feature vector *i*. The splat designates convolution operator. *B* is a fixed *base image. T*_*i*_ is the convolution of the base image and a kernel formed from the input feature vector *i*.

In the test (inference) phase, each input vector is converted to image using the same procedure used in training. The resulting image is then classified using the previously trained CNN.

We used simple accuracy (percentage of correctly classified samples) to compare the classification performance.

To establish non-CNN results, we used a custom ML program for classifying infectious diseases [4], implementing XGBoost, LightGBM, SVM with linear and RBF kernel, multi-layer perceptron (MLP) and regularized logistic regression. XGBoost, LightGBM, logistic regression and MLP used random search [5] over hyperparameters supported by the scikit-learn [6] python module and Tensorflow (MLP). SVM classifiers used grid search. The hyperparameter configurations were based on previous experience in this application domain. The analyses proceeded as follows:

- for each hyperparameter configuration (HC), estimate accuracy using 5-fold cross-validation
- select HC with highest accuracy
- train model on entire training set
- apply the model to validation set and report accuracy

In addition, we used h2o.ai Driverless AI software [7] to find the best model. The Driverless AI run used maximum accuracy, maximum time and minimum interpretability settings, to ensure fair comparison with other sofware. All other settings were the defaults.

We reported the highest validation set accuracy as the non-CNN result.

We did not attempt hyperparameter tuning of the ResNet, except minor change to the weight decay parameter. The architecture (ResNet-34), hyperparameters and the software were adopted from the fast.ai course [8].

All computations were performed on a 20-core computer. CNN training was performed using fast.ai library [8] running on NVIDIA Titan X GPU.

### 2.1 Materials

To assess performance of TAC, we applied it to the problem of classifying blood samples from patients with bacterial or viral infections. Accurate diagnosis of infection type could help guide the use of antibiotics and improve management of sepsis, a life-threatening condition. The process of generating the data is described previously [4]. Briefly, blood samples were processed using a gene expression instrument (gene chip microarrays or Nanostring instrument), normalized and pre-processed. The result was a set of 2331 training samples and 259 validation samples. Collectively, these 2,590 samples present clinical, biological and technical heterogeneity observed in the real world as the come from 41 independent studies profiled using microarrays or Nanostring. We split the samples by the gene expression processing platform to demonstrate *platform transfer*: the training samples were assayed using predominantly (90%) gene expression microarrays, while the validation samples were assayed using a Nanostring gene expression instrument.

Partition by type was used to demonstrate the platform transfer: learning on one platform and testing on another. The proportions of bacterial infections were 52.9% in the training set and 40.9% in the validation set.

## 3 Results

We trained and evaluated the classifiers using the training and validation sets described in the Section 2.

The initial dataset contained 29 input genes that were chosen based on prior research in diagnosing infections using host immune response to pathogens [9, 10, 11]. Because the nearest odd square to 29 is 25, we decided to remove 4 genes with smallest variance, estimated using the training data. The remaining 25 genes were used to form the 5×5 image filter kernels for TAC.

We used several base images: MNIST digits, black-and-white rose image, uniform random noise base image, and an image which contains combination of random noise and labeled axes, referred to as combined image. The MNIST digits did not perform well, perhaps due to low source resolution, and were abandoned (data not shown). The transformed images were resized to 400 by 400 pixels. The Fig. 2 shows an effect of the convolution between the rose base image and gene expression vectors.

**Figure 2:**
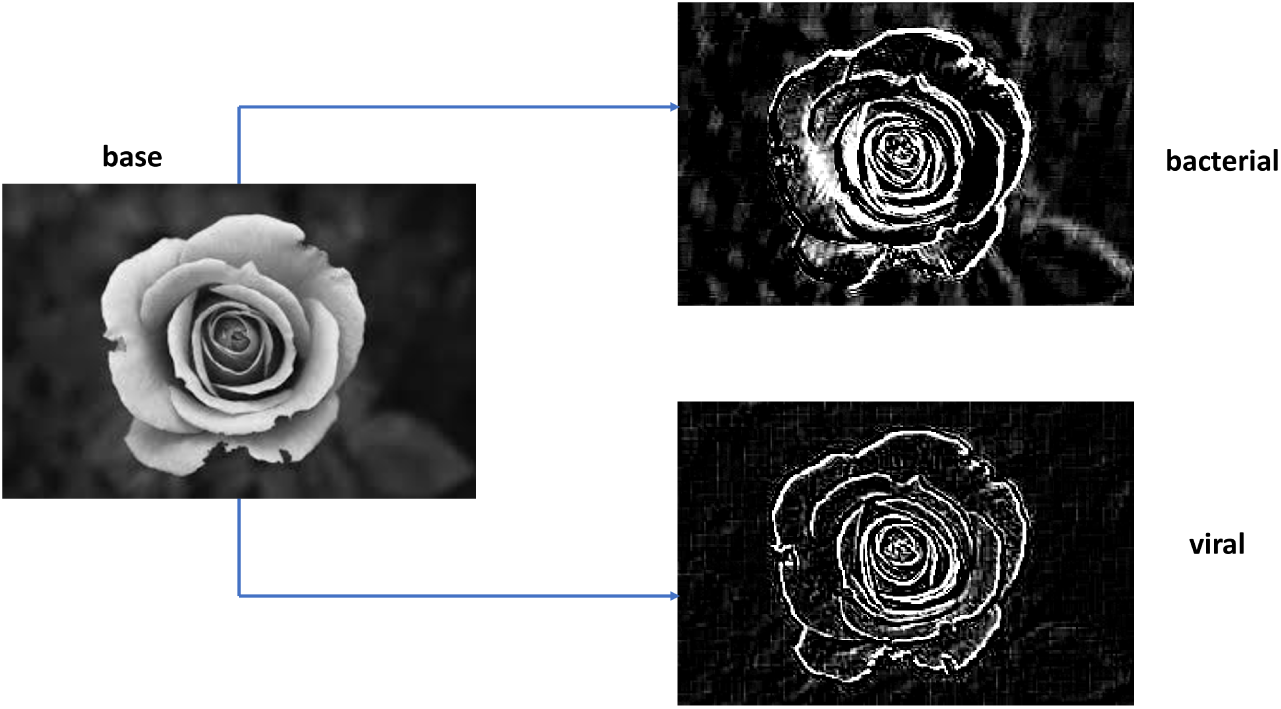
The core of the TAC approach: convolution between a base image and input feature vectors. The figure shows rose base image and corresponding filtered images. The filtered images are obtained by convolving the base image with filters formed from representative bacterial and viral feature vectors. See Section 2 and Fig. 1 for details.

Among the non-CNN classifiers, linear Support Vector Machine achieved the highest accuracy (89.6%) with 100 hyperparameter configurations, whereas multi-layer perceptron (MLP) had the lowest (81.5%), despite taking the longest time to train (Table 1).

**Table 1:**
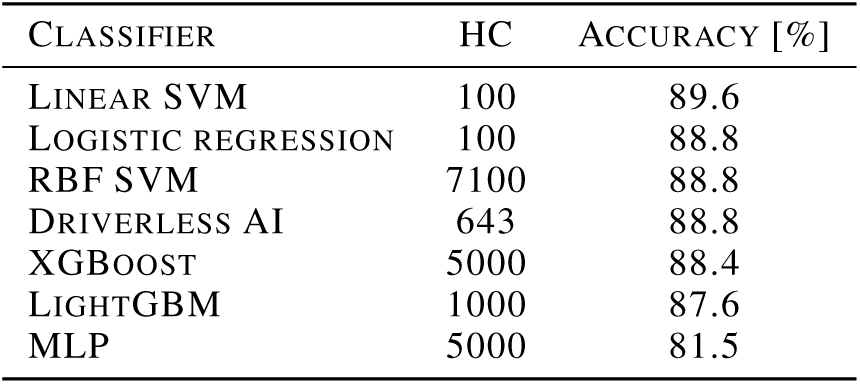
Hyperparameter configurations and classification accuracies for non-CNN classifiers. The HC column contains number of hyperparameter configurations searched.

| CLASSIFIER | HC | ACCURACY [%] |
| --- | --- | --- |
| LINEAR SVM | 100 | 89.6 |
| LOGISTIC REGRESSION | 100 | 88.8 |
| RBFSVM | 7100 | 88.8 |
| DRIVERLESS AI | 643 | 88.8 |
| XGBOOST | 5000 | 88.4 |
| LIGHTGBM | 1000 | 87.6 |
| MLP | 5000 | 81.5 |

In comparison, TAC achieved the same accuracy as linear SVM in 25 epochs, where each epoch took approximately 30 seconds (Fig. 3).

**Figure 3:**
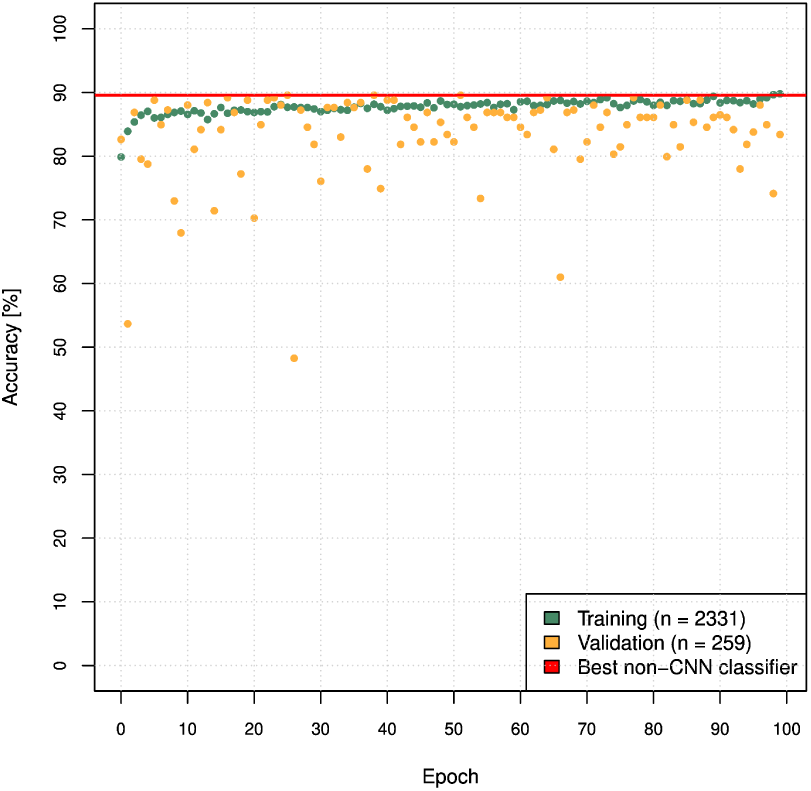
ResNet training using TAC-transformed rose image.

Next, we trained the network for a large number of epochs (> 10000) using several base images. The maximum recorded accuracy was 91.1%, though this result could be due to overfitting over many epochs (Fig. 4). Interestingly, this result was obtained using the combined image. The best accuracy achieved with purely random noise image was 90.0%.

**Figure 4:**
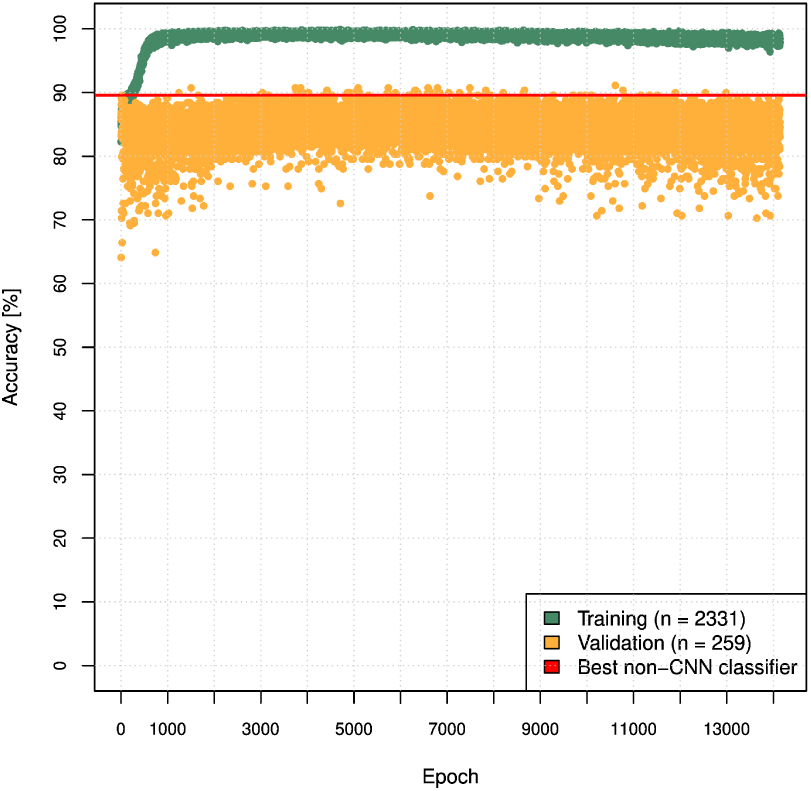
ResNet training using TAC-transformed combined base image (random noise and labeled axes), performed for ∼ 14000 epochs.

The results are summarized in Table 2.

**Table 2:** Classification accuracies for the best non-CNN classifier and TAC with different base images and epochs. Best: highest accuracy achieved over the given number of epochs along with 95% confidence interval. Avg.: average accuracy over the epochs.

| CLASSIFIER | BASE IMG. | EPOCHS | BEST | AVG. |
| --- | --- | --- | --- | --- |
| NON-CNN | | | 89.6± 3.7 | |
| TAC | RANDOM | 50 | 89.2± 3.8 | 77.4 |
| TAC | RANDOM | 3207 | 90.0± 3.7 | 77.8 |
| TAC | ROSE | 50 | 89.6± 3.7 | 82.5 |
| TAC | ROSE | 18418 | 91.1± 3.5 | 84.0 |
| TAC | COMBINED | 50 | 89.6± 3.7 | 83.8 |
| TAC | COMBINED | 14146 | 91.1± 3.5 | 84.6 |

The energy consumption for the TAC training was approximately 126kWh during about three weeks of research. We are unable to report energy consumption for the ML analyses because we used software libraries for which energy tracking is not user-accessible.

## 4 Discussion

We propose TAC for applying CNN to tabular data. TAC uses tabular data to create image filters that are applied to a fixed base image. To the best of our knowledge, this approach is novel, and our study is the first one to classify clinically relevant tabular data using CNNs. Our results show that the TAC approach can match accuracy of classifying gene expression data compared with non-CNN machine learning classifiers.

One could contemplate that neural networks are successful in classifying images, sound and text because such data sources have internal data structure. For example, in images, the internal data structures are various shapes at various levels of complexity, which are detected by neural network layers. In contrast, such structures are thought lacking in tabular data, and therefore neural networks do not achieve superior performance. Alternatively, tabular data may possess internal data structure that is not detectable by human senses. If this is correct, perhaps the convolutional filtering of tabular data proposed in this paper translates those structures into image structures and enables their subsequent discovery using CNNs.

In our experiments, the impact of input base image was somewhat inconclusive. We slightly surpassed non-CNN classifiers using several different base images, including a random noise image. This is surprising because there are no discernible visual features in the transformed random noise images and thus it is not clear why CNNs would be beneficial. Nevertheless, both overall and average accuracy for a random noise image were inferior to “real” base images, which should therefore preferably be used for TAC. In addition, the convergence on real images could potentially be sped up by applying transfer learning. Since random images do not have recognizable features, it is unlikely they would benefit from it. Overall the best result was achieved using “combined” base image consisting of noise and digits. Deeper understanding of impact and selection of base images is an area of active research for us.

One possibility of further improving results is to use a small set of base images instead of a single one. Each filter would be applied to all base images in the set, and the images are combined into a single image, which corresponds to the input feature vector. This could potentially improve the discrimination among the TAC input images. This is left for future research.

The order of the 25 genes used to create convolutional filters was arbitrary. It produced visible patterns (Fig. 2) in the filtered images, which enabled accurate classification of the images using CNN. One way to potentially improve accuracy might be to reorder the genes and/or generate multiple filtered images for each input feature vector (one per specific ordering of genes), and then combine those multiple images for classification by CNNs. The rationale is that different orderings might generate different patterns in filtered images, whose combination might improve discrimination.

We used ResNet architecture implementation as described in the fast.ai course. In some problems, the results could perhaps be improved by considering more advanced architectures. However, in the gene expression dataset that we used that would probably not be necessary because the ResNet already achieved 100% training classification accuracy, and the most promising path forward is probably regularization.

We used accuracy as performance metric because it is adequate for assessing the potential of TAC. However, it is not a relevant metric for our target application (classification of infections) due to asymmetric costs, and in the future we plan to assess TAC using clinically-relevant metrics.

TAC achieved absolute best accuracy (point-estimate) of 91.1%. The best accuracy achieved using non-CNN was 89.6%. This means that TAC reduced the fraction of incorrect diagnoses (bacterial or viral) by about 14%, which could be clinically significant. However, this result could be optimistically biased because it required several thousand epochs of CNN training. Using 50 epochs, where overfitting is much less likely, we achieved similar performance as the non-CNN classifiers.

The confidence intervals for the point estimates overlap because a relatively small validation set was available. Further research with larger datasets is in progress to confirm or refute the findings.

Comparison of classifier performance is a complex topic, in particular between deep learning classifiers and classical ML because they use, traditionally, different performance assessment methods. We used the best practices known to us. For non-neural-network approaches, one typically searches for best hyperparameter configuration using cross-validation [12] or similar model selection methods, then applies the best model to the validation set. Thus, the validation set is generally “seen” a handful of times before the final model is locked. In contrast, in CNN practice a trained network is applied to the validation set after each epoch, which leads to potentially excessive use of the validation data. This is not a significant problem in large sample size scenarios like ImageNet, but it is a challenge in clinical applications which typically have far fewer samples. One way to approach this would be to use cross-validation with CNNs, but the computational complexity might be a limiting factor. This is an open question left for future research.

The computational complexity of TAC is essentially equivalent to complexity of CNNs. The kernel transformation and convolution are performed once for each dataset and take comparatively negligible amount of time during training. At inference time, the transformation of input vector to an image amounts to another convolution, which is a negligible increase in complexity of today’s state-of-the-art architectures.

We did not apply transfer learning in this paper, in part due to limited resources. We hypothesize that it could improve convergence of the best models, in the case of realistic base images. In the case of random base images, it is unlikely that transfer learning would make a difference because there aren’t any visually discernible features to transfer.

The TAC approach could be nearly fully automated if off-the-shelf CNN and hyperparameters are used, and thus the classification task could be significantly simplified compared with non-CNN ML, which involves hyperparameter tuning, feature engineering etc.

One limitation of our method is that it applies to relatively low-dimensional input spaces, in order to construct reasonable-sized filters. To generalize to higher-dimensional problems, several approaches could be considered. For example, the feature vectors could be split in segments of length 25 or 49, each yielding 5×5 or 7×7 kernel. Each kernel could be applied to the base image, and then images could be combined together into a single image which is then processed using CNNs as already described in the paper. Alternatively, the images could be stacked as input channels. This still would not work for extraordinarily high numbers of features, such as those present in some genomic datasets. In those cases, some form of initial feature (gene) selection would be required prior to applying TAC. This is commonly done even with classic ML analyses of such datasets.

We analyzed single, albeit important, dataset. Further research with additional datasets is in progress.

In this paper we analyzed continuous data. For categorical data, a common approach one could apply is entity embeddings [13]. Once embedded, data could be used in TAC directly.

In conclusion, our results suggest an alternative and promising path (TAC) for classification of tabular data using CNNs.

